# Neural correlates of perisaccadic visual mislocalization in extrastriate cortex

**DOI:** 10.1101/2023.11.06.565871

**Authors:** Geyu Weng, Amir Akbarian, Kelsey Clark, Behrad Noudoost, Neda Nategh

## Abstract

When interacting with the visual world using saccadic eye movements (saccades), the perceived location of visual stimuli becomes biased, a phenomenon called perisaccadic mislocalization, which is indeed an exemplar of the brain’s dynamic representation of the visual world. However, the neural mechanism underlying this altered visuospatial perception and its potential link to other perisaccadic perceptual phenomena have not been established. Using a combined experimental and computational approach, we were able to quantify spatial bias around the saccade target (ST) based on the perisaccadic dynamics of extrastriate spatiotemporal sensitivity captured by statistical models. This approach could predict the perisaccadic spatial bias around the ST, consistent with the psychophysical studies, and revealed the precise neuronal response components underlying representational bias. These findings also established the crucial role of response remapping toward ST representation for neurons with receptive fields far from the ST in driving the ST spatial bias. Moreover, we showed that, by allocating more resources for visual target representation, visual areas enhance their representation of the ST location, even at the expense of transient distortions in spatial representation. This potential neural basis for perisaccadic ST representation, also supports a general role for extrastriate neurons in creating the perception of stimulus location.

## Introduction

Saccades are rapid eye movements that shift the center of gaze to a new location in the visual field. Changes in visual perception occur around the time of saccades^1,2^. For example, our subjective experience of the visual scene remains stable across the abrupt changes of the retinal image during saccades. This phenomenon is called visual stability, and many studies have attempted to explain the mechanism behind it^3^. Several other perceptual phenomena which occur around the time of saccades have also been studied psychophysically. For example, there is a general reduction in visual sensitivity during saccades, a phenomenon called saccadic suppression or saccadic omission, that has been reported in both macaques and humans^4–7^. Saccadic eye movements also alter our perception of time^8^. Another phenomenon is perisaccadic mislocalization, in which the perceived location of visual stimuli appearing near the time of a saccade is biased. Perisaccadic mislocalization was first discovered as a perisaccadic shift, a unidirectional mislocalization parallel to the saccade, when the experiments were done in darkness with human subjects^9–12^. Later studies have demonstrated perisaccadic compression ^13,14^, which is mislocalization towards the saccade target (ST), when the subjects make saccades with background illumination and visual references^15–18^.

Perisaccadic visual perception in macaques is qualitatively similar to humans^4^, and many studies have investigated the neurophysiology of perisaccadic visual perception in nonhuman primates^19–26^. Some neurons in the extrastriate visual areas and prefrontal cortex show a sensitivity shift to the postsaccadic receptive field (RF) even before the saccade, a phenomenon often referred as future field remapping^27,28^. There is also another phenomenon, called ST remapping, in which neural RFs shift towards the ST around a saccade^29–38^. Both future field and ST remapping can be observed in the same experiments in the same group of neurons^39–42^. It has been suggested that the RF remapping is associated with perisaccadic mislocalization^19,43,44^, and some studies have used computational approaches to predict perisaccadic perception of space based on neural responses^45–48^. Although these studies have generated insightful experiments, theories, and hypotheses, they usually start with assumptions about the function of visual areas or have a limited precision in accounting for the time-varying relationship between neural modulations and perceptual alterations on the millisecond timescale of saccades. By quantifying the statistical dependencies of spiking responses on several behavioral (e.g., eye movement) or external (e.g., visual stimuli) variables, however, point process statistical models provide a powerful means to capture the encoding and decoding of visual information as continuously varying with eye movements, with no assumption on the function of neurons. To investigate the neural basis of perisaccadic mislocalization, this study used a time-varying generalized linear model framework capable of capturing the fast spatiotemporal dynamics of neural sensitivity around the time of saccades^42,49^, and examine the link between perisaccadic visual responses and visuospatial perception.

In this study, we used a combined experimental and computational approach built upon neuronal responses in the middle temporal (MT) cortex and area V4 of rhesus macaque monkeys. We first assessed each neuron’s sensitivity to each location of visual space across time relative to the saccade (neuron’s kernels) using a statistical model fitted on the recorded spiking data during a visually guided saccade task with visual stimulation. We quantified the representational spatial bias using the spatiotemporal kernels of populations of neurons, based on the similarity in neural sensitivity to neighboring probe locations, without assumptions about the downstream readout mechanisms. We then used this measure of spatial bias to identify the perisaccadic changes in sensitivity which drive it, and linked them to neural responses.

We found that neurons with RFs close to the ST do not contribute to spatial bias. In contrast, perisaccadic spatial bias in the direction opposite to the saccade vector can be accounted for by neurons with RFs farther from the ST. These neurons showed perisaccadic and postsaccadic sensitivity changes near the ST (a.k.a. ST remapping) that contributed to spatial bias. We found unexpectedly that the time course and response components of the spatial bias matched that of another perisaccadic perceptual phenomenon, namely the enhancement of neural sensitivity around the ST. This representational ST enhancement can link to presaccadic enhanced ST perception reported in psychophysical studies^35,37,50^ and to presaccadic increased stimulus selectivity^24,26,51,52^ or ST remapping^38,39^ evidenced in neurophysiological studies. The shared neural response components underlying the ST representational enhancement and bias suggest that the brain likely trades off and prioritizes saccade target representation with consequential biases in location perception.

Taken together, our findings highlight a potential neural basis for perisaccadic mislocalization, supporting a role for extrastriate neurons in the perception of stimulus location and linking ST remapping to perisaccadic spatial bias with simultaneous representational enhancement of the ST area.

## Results

To examine the neural basis of perisaccadic spatial biases in perception, we recorded the responses of extrastriate neurons (see Methods). We analyzed the activity of 300 neurons from MT and 147 neurons from area V4, recorded while monkeys performed a visually-guided saccade task (Fig. 1a). During the first fixation period, the monkey first fixated on a fixation point (FP), and a ST appeared 13 degrees of visual angle (dva) away either to the left or the right horizontally, while the monkey held the fixation. When the FP disappeared, the monkey had to make a saccade to the ST and maintain fixation on the ST during the second fixation period. A series of probe stimuli were presented throughout the task while the monkey fixated and made a saccade. Only one stimulus was presented at a time, selected from a 9×9 grid of possible locations, and each stimulus appeared for 7 ms. The probe grid was adjusted to cover the FP, ST, and estimated RF of the neuron. In order to computationally investigate the mechanism of spatial bias, we developed an encoding model which quantitatively characterizes the neuron’s input-output relationship and captures the neuron’s sensitivity map with high temporal precision throughout the eye movement task (see Methods). The model traces the time-varying dynamics of a neuron’s sensitivity across saccades with high-dimensional spatiotemporal kernels. For each of the 81 probe locations, we decomposed all times of the neural response relative to saccade onset and delays (times of the stimulus relative to each response time) into 7-ms bins to form 4-dimensional spatiotemporal units (STUs). We define each spatiotemporal unit (STU) based on the response of the neuron to one probe location at each time and delay bin (Fig. 1b). Each STU is assigned a single numerical weight (STU weight) after fitting the model to the spiking data. The combinations of these STUs across time and delay values constitute the neuron’s spatiotemporal kernel map at each probe location. Figure 1c shows the STU weights at an example probe location around the RF of an example neuron across time and delay. When responding to a probe stimulus at that location, the STUs comprise kernels that represent how a neuron’s sensitivity changes across time from saccade onset and across delays. We used the Sparse Variable Generalized Linear Model (SVGLM)^53^ to estimate the STU weights and the resulting kernels by fitting to the neuron’s spiking responses (see Methods and Supplementary Fig. 1). A signal representing the stimulus across the 9×9 grid locations passes through spatiotemporal kernels representative of the neuron’s time-varying spatiotemporal sensitivity and added to the time-varying baseline neural activity relative to saccade onset captured by an offset kernel, and the feedback signal generated using a post-spike filter representing the effects of spiking history. The combined signal is then passed through a sigmoidal nonlinearity capturing the spike generation. The resulting firing rates are used to generate spikes with the Poisson spike generator. These spikes are then fed back to the circuit through the abovementioned post-spike history (Supplementary Fig. 1a). All the model components are learned via an optimization process to directly estimate the recorded spiking activity (see [49] for details). The kernels estimated from the model reflect the temporal sensitivity of the neuron at each probe location (Fig. 1c).

**Fig. 1.**
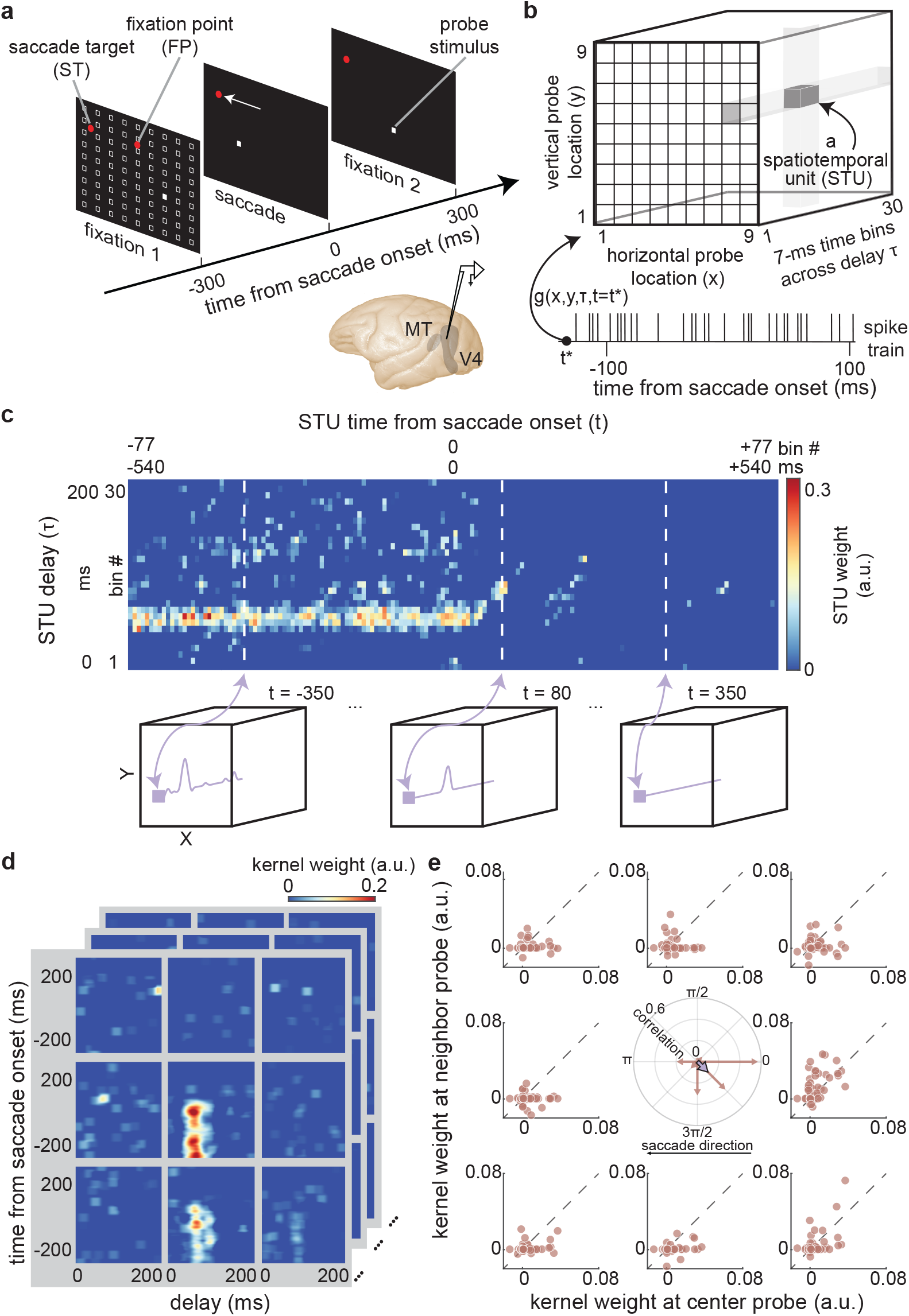
Experimental and computational paradigm for measuring spatial bias. **a.** Schematic of the visually-guided saccade task with probes. Monkeys fixate on a central fixation point (FP), then a saccade target (ST) point appears in either horizontal direction. After a randomized time-interval (700:1100 ms), the FP disappears, cueing the monkeys to saccade to the ST. Throughout the task, a series of pseudorandomly located probes appear in a 9×9 grid of possible locations (white squares). Only one probe is on the screen at each time, for 7 ms. Neurons were recorded from the middle temporal (MT) area or area V4. **b.** Composition of the neuron’s sensitivity map for an example timepoint relative to saccade onset using spatiotemporal units (STUs) across locations and delay bins of 7-ms. **c.** STU weights across time and delay characterizing the sensitivity dynamics of a sample neuron for an example probe location. At each of the 9×9 locations, the neuron’s sensitivity is captured by kernels that comprise the weighted combination of STUs and represent the time-varying spatiotemporal sensitivity of the neuron across time and delay. Each kernel has spatial (x, y), time (t), and delay (𝜏𝜏) dimensions, and the spatial coordinates are based on locations on the screen. Time refers to the time of response relative to saccade onset (-540:540 ms), and delay refers to the time of stimulus relative to particular response time (response 0:200 ms after stimulus onset), discretized in bins of 7-ms. **d.** Each layer represents the spatiotemporal kernels of one neuron at 9 probe locations around the neuron’s RF during the initial fixation. These kernels are calculated for each neuron in each ensemble (z dimension in the panel). **e.** Scatter plots show the kernel weights for the center probe closest to the ST vs. those for the eight surrounding locations, for each neuron in an example ensemble (n = 53 neurons), for a particular time and delay combination (time = 100 ms and delay = 110 ms). Eight correlation vectors can be computed, using correlation strengths as magnitudes and the relative probe positions as directions (brown arrows in center panel). The eight vectors are averaged to a single vector (purple arrow in center panel) which represent possible bias in reading out the location of that center probe from the population sensitivity.

Next, we developed a procedure to measure spatial bias based on the neurons’ estimated kernels. We made the assumption that similar neural sensitivity to probes appearing at neighboring locations could create uncertainty in a readout of the stimulus location by a downstream area, which would lead to a bias in spatial perception. In other words, if during a saccade, the population response to one probe becomes similar to that of a neighboring probe, we can assume a representational bias toward that neighboring location, without specifically modeling downstream readout mechanisms. To examine the neural basis of spatial bias, we analyzed the similarity between the spatiotemporal kernels at pairs of probe locations in a population of neurons. For the sensitivity analysis at the population level, we divided the neurons into ensembles based on their RF locations. Neurons recorded with the same RF, ST, and grid arrangements were grouped as an ensemble, and each ensemble had a minimum of 10 neurons. Figure 1d shows the kernel maps at 9 probe locations around the RF for an example ensemble of neurons. For each ensemble, we measured the similarity between the neural sensitivity at neighboring locations by taking cosine correlations between the kernel weights at the center probe and the kernel weights at each of its eight neighboring probes; these correlations are calculated at each time and delay value. For the rest of the paper, correlation always refers to cosine similarity between the kernel weights for neighboring probes across neurons in an ensemble. In this study, we focused on examining the spatial bias around the ST because prior psychophysics often reported perisaccadic mislocalization close to the ST. Next, we define a spatial bias measure based on ensemble sensitivity to probes near the ST. Figure 1e shows the correlations of kernel weights between an example probe close to the ST and its 8 neighboring probes, for an example ensemble of 53 neurons at time 100 ms and delay 110 ms. The correlation coefficient between the central probe and its neighboring probes (central polar plot) indicates the similarity of neural sensitivity in each direction at that central probe location (Fig. 1e, brown arrows). We then averaged over the eight vectors at each probe location to get one vector (Fig. 1e purple arrow). Since the saccade direction was either to the left or the right horizontally, we focused on the horizontal projections of the average vectors, which we defined as the spatial bias. Values were normalized according to saccade direction so that positive always means the same direction as the saccade, and negative means the opposite direction from the saccade. This spatial bias measurement allowed us to predict potential mislocalization of stimuli based on the kernels of the SVGLM fit to neural data.

Next, we examined how spatial bias changed over time relative to the saccade and the stimulus. Each kernel map has its own time and delay dimensions, so we measured spatial bias maps across time and delay for each of the 7×7 probe locations for each ensemble. Figure 2a shows the spatial bias over delay, at time 100 ms, at a probe location close to the ST for an example ensemble, and Figure 2b shows the spatial bias over time at delay 110 ms for the same ensemble and probe location. We normalized the spatial bias so that each ensemble has values ranging from -1 to 1. In this study, we focused on the probe locations around the ST. For each ensemble, we selected 6 locations around the ST and averaged their bias maps. We then averaged the bias maps for 15 ensembles (447 neurons) (Fig. 2c). Taking the mean over delays of 50:100 ms, we observed a negative bias (which means a significant bias in the direction opposite to the saccade direction) of -0.13±0.06 around the ST for ∼50:150 ms after saccade onset (Fig. 2d). To find out whether the amount of bias correlates with the eccentricity of neurons’ RF location relative to the ST location, we grouped ensembles of neurons based on *d* – the distance between the RF center and the ST (Fig. 2e). Ensembles with *d* < 11 dva showed very little bias compared to other groups (0.02±0.13). To examine the variability of neurons within each ensemble and its possible effect on the amount of bias, we resampled 90% of the neurons in each ensemble to compute 100 samples of spatial bias for each ensemble. The mean spatial bias in the perisaccadic window of 50:150 ms demonstrates that most of the ensembles with RFs closer to the ST show less spatial bias, and the standard error of the mean shows that the phenomenon within each ensemble is consistent (Fig. 2f). Thus, our population of neurons showed perisaccadic spatial bias opposite to the saccade direction, primarily driven by neurons with RFs far from the ST.

**Fig. 2.**
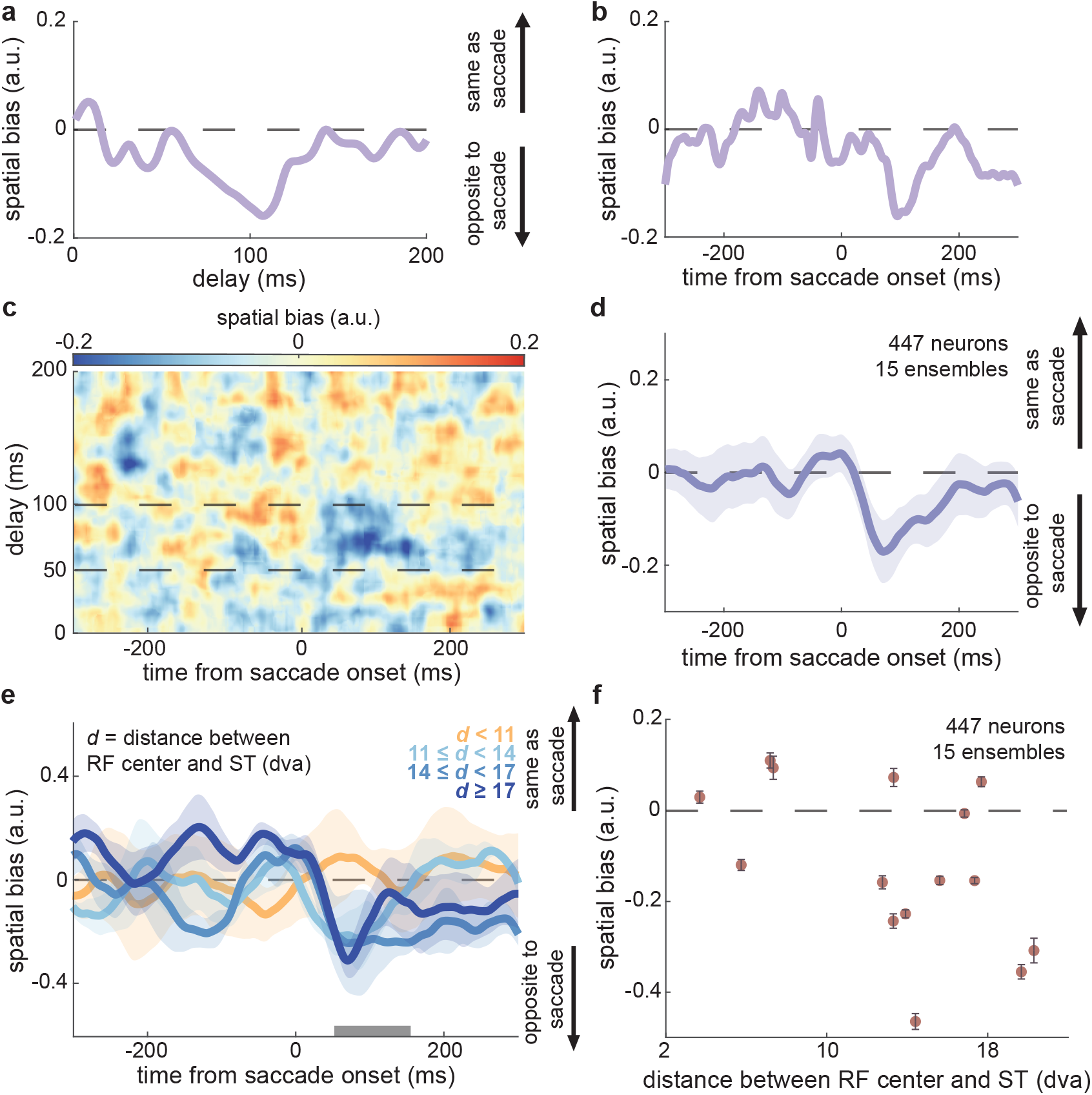
Quantifying the spatial bias and its dynamics over time, delay, and ensembles. **a.** The spatial bias as a function of stimulus delay values, for a probe that appears at a location close to the ST, measured using the neurons’ sensitivities at time 100 ms after saccade onset in an example ensemble. **b.** The spatial bias as a function of time relative to saccade onset, for the same probe and ensemble in (a), measured using the neurons’ sensitivities at delay 110 ms relative to each timepoint (x-axis). **c.** Mean spatial bias across time and delay, for 6 probe locations around the ST, averaged across all 15 ensembles (n = 447 neurons). Dashed lines indicate delay values used in (d). **d.** Mean spatial bias over time, for delay 50:100ms, for 6 probe locations around the ST, for all 15 ensembles. Shaded area represents the standard error of the mean (SEM) across ensembles. **e.** Spatial bias over time from saccade onset, for ensembles with various distances between their neurons’ RF center and the ST (*d*). There are 4 ensembles with *d* < 11, 4 ensembles with 11 ≤ *d* < 14, 3 ensembles with 14 ≤ *d* < 17, and 4 ensembles with *d* ≥ 17. Shaded area represents SEM across ensembles. **f.** Spatial bias for the 15 ensembles during the perisaccadic window (50:150 ms from saccade onset, gray bar in (e)), plotted against the distance between RF center and the ST for each ensemble. Error bars indicate the SEM of the bias estimate over resampling the neuronal population in each ensemble (n = 100 samples, 90% of neurons in each sample).

The above results show that the perisaccadic changes in the spatiotemporal sensitivity of MT and V4 neurons could account for changes in spatial perception during eye movements, but so far, we have focused on the representation at the population level and model-based neural sensitivity measurements. In order to find out which components of the neuronal response of which neurons account for the perisaccadic alteration in the readout of location, we used an unsupervised approach to search for response components that are specifically related to spatial bias. In this study, spatial bias was defined based on similarity in the population representation of neighboring probe stimuli captured by the neurons’ spatiotemporal kernels; since the kernels are comprised of STUs, manipulation of certain STUs can change the kernels and thereby affect the similarity between the population sensitivity at neighboring locations. This assumption-free alteration in the model enables us to determine which of the modulated STUs are necessary for creating spatial bias. Based on this rationale, we defined a bias index according to the difference between the center kernel and each neighboring kernel across times and delays. Nulling each modulated STU one by one we can quantify their effect on the kernel similarity using this bias index, and systematically identify the bias-relevant STUs (see Methods; Supplementary Fig. 2). Using this unbiased search in the space of STUs, we found different phenomena for ensembles with different distances between the RF center and the ST (*d*), so we divided the ensembles into two groups (*d* < 11 dva and *d* ≥ 11 dva) to examine their bias-relevant STUs separately (Fig. 3a). For ensembles with *d* < 11 dva, there was a set of bias-relevant STUs around time 60:90 ms and delay 80:110 ms. For ensembles with *d* ≥ 11, there were two areas of bias-relevant STUs – one around time 60:100 ms and delay 60:110 ms, and the other one around time 110:280 ms and delay 50:100 ms. After removing all the identified bias-relevant STUs, we recomputed the spatial bias over time, and the previously observed bias around 50:150 ms after saccade onset was significantly reduced (Fig. 3b; -0.03 ± 0.04, *p* = 0.04), confirming that the identified set of STUs drive this bias. Thus, by leveraging the capabilities of the model to decompose the spatiotemporal sensitivity of individual neurons, we were able to identify the specific changes in neural sensitivity that contribute to perisaccadic spatial bias.

**Fig. 3.**
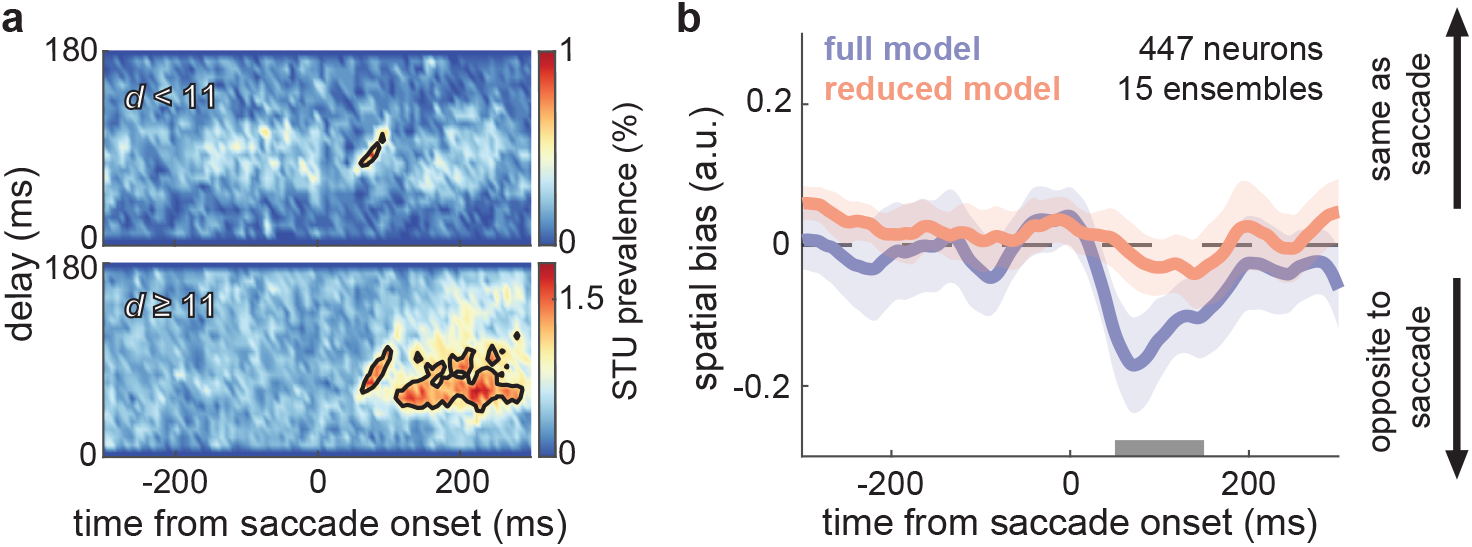
Identifying and validating bias-relevant sensitivity components. **a.** Prevalence of bias-relevant STUs, over delay and time from saccade, for ensembles with *d* < 11 dva (top, n = 4 ensembles) and the other with *d* ≥ 11 dva (bottom, n = 11 ensembles). The black contours show the outline of STUs above 60% of the maximum prevalence. **b.** Mean spatial bias over time from saccade onset, in the full model (purple), and in the reduced model (pink) where all the bias-relevant STUs associated with the ST probes and their neighbor probes are removed. Plots show mean±SEM across 15 ensembles confirming that the spatial bias is significantly reduced for the 50:150 ms time window after saccade onset (gray bar) (*p* = 0.04).

To interpret how the saccade-related changes in STUs link neurophysiological activity to a biased readout of location information, we wanted to relate them back to the neural responses. The model allows us to generate responses to synthetic stimuli, and compare the predicted neural response during fixation and perisaccadically. We first examined the model-predicted response for ensembles with *d* < 11, and transformed the time and delay of the bias-relevant STUs to a stimulus-aligned response (Fig. 4a). To investigate the neural response underlying the perisaccadic change in spatial bias, we looked at how neurons responded to probes on different sides of the ST. Data from ensembles recorded with leftward saccades have been flipped to be combined with those recorded with rightward saccades (Fig. 4b). Out of the six probes around the ST, we called the three probes closer to the FP the “near” probes and the other three probes the “far” probes (Fig. 4b). RFs of neurons in ensembles with *d* < 11 mostly cluster between FP and ST (see prevalence in Fig. 4b), resulting in the near probes falling close to the RFs. Based on figure 4a, we averaged the model response for near vs. far probes over fixation (-500:-100 ms) and perisaccadic (-20:10 ms) windows, and used the neurons’ responses from experimental recordings as validation (Fig. 4c). We specifically compared the responses in 60-ms windows of time from stimulus onset, around the peak of the fixation and perisaccadic responses (fixation: 50:110 ms, perisaccadic: 70:130 ms). During fixation, there was a greater model-predicted response to near probes vs. far probes (near = 1.05±0.01, far = 1.01±0.01, *p* < 0.001, n = 94 neurons), consistent with the near probes being closer to the RF centers. There was an increase of model-predicted response perisaccadically for both near and far probes, but more of an increase for far probes, such that the perisaccadic response ended up being similar for near and far probes (near = 1.05±0.02, far = 1.07±0.02, *p* = 0.13, n = 94). The model closely predicted the response from actual neurons during both the fixation window (experimental values: near = 1.04±0.01, far = 0.94±0.01) and the perisaccadic window (experimental values: near = 1.20±0.04, far = 1.20±0.04). We measured the statistical difference between the actual firing rates of neurons in response to near vs. far probes (Fig. 4d), in 60-ms windows matched to their evoked responses. During the fixation period, from 50 to 110 ms after stimulus onset, the firing rate evoked by near probes was significantly higher than that for far probes (near = 38.57±2.30 Hz, far = 35.21±2.18 Hz, *p* < 0.001). During the perisaccadic period, from 70 to 130 ms after stimulus onset, there was no statistically significant difference between the firing rates in response to near vs. far probes (near = 43.39±2.63 Hz, far = 43.13±2.57 Hz, *p* = 0.50). These neural responses show that neurons with RFs close to ST responded more to near probes during fixation, but responded equally to both near and far probes around the time of saccades. The lack of difference in response indicates that there is no neural bias towards either side of the ST around the time of eye movements, which explains the absence of spatial bias for ensembles with *d* < 11.

**Fig. 4.**
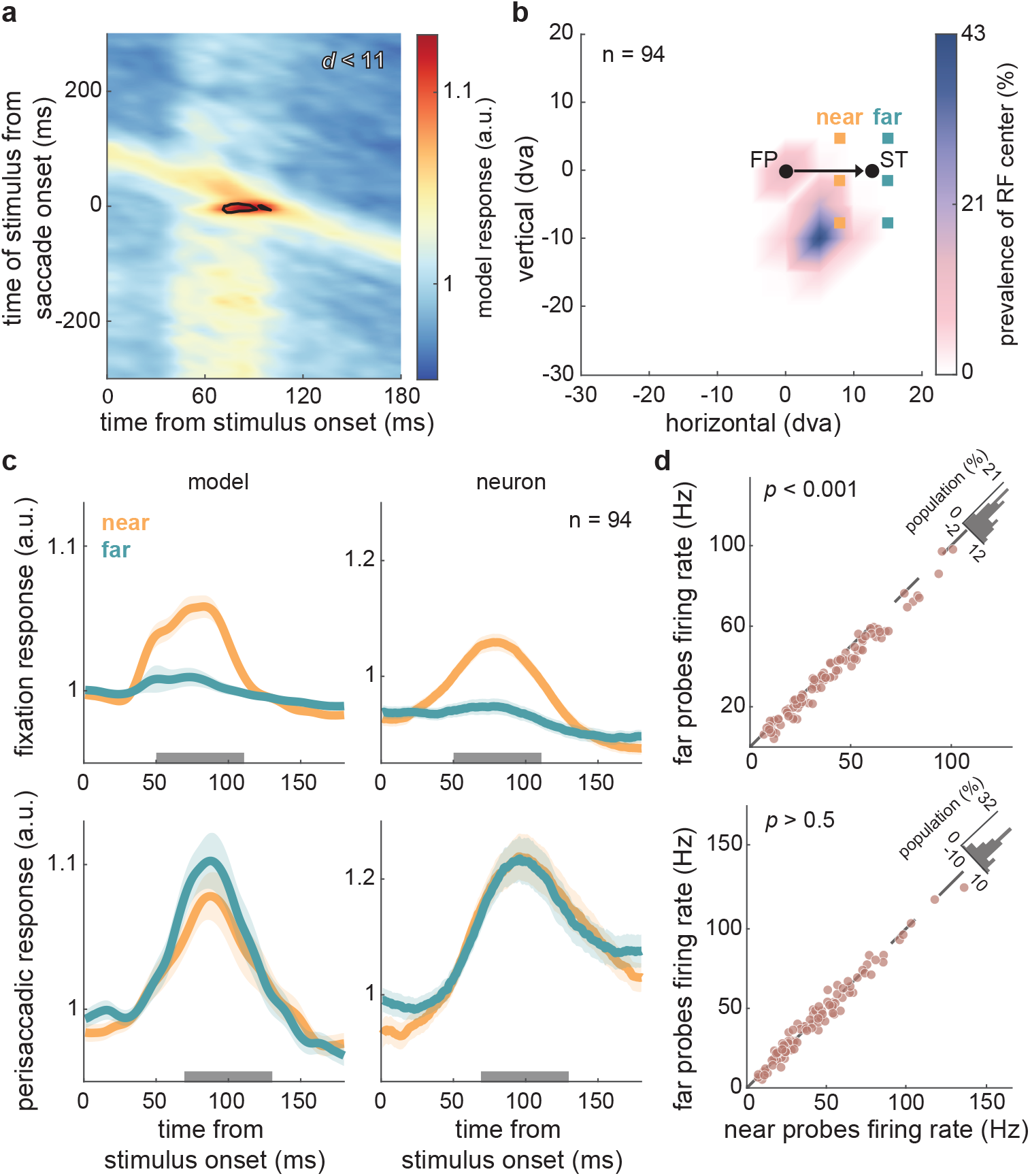
Identifying the neural correlates of spatial bias around the ST area from neurons whose RFs are located near the ST. **a.** Bias-relevant STUs and model-predicted response, plotted as a function of time of stimulus from saccade onset (y-axis) and time of response from stimulus onset (x-axis). Shown for ensembles with *d* < 11 dva (n = 4 ensembles, 94 neurons). **b.** Map of RF centers relative to the FP and ST using all ensembles used in (a), and the probe locations around the ST. Prevalence of RF center (colorbar) indicates the percentage of neurons with RF centers in the corresponding location. “Near” probes are the three probes on the side of the ST towards the FP (orange), while the “far” probes are on the other side of the ST (green). **c.** Mean of normalized model-predicted responses (left) and actual neural responses (right) over time from probe onset, for near (orange) and far (green) probes, during fixation (top; -500:-100 ms) and perisaccadic (bottom; -20:10 ms) windows. Mean±SEM across models or neurons in the ensembles; gray bars show analysis windows used in (d). **d.** Comparison of actual neural responses to near vs. far probes (n = 94 neurons), for the fixation period (top, *p* = 5.38e-12) and perisaccadic period (bottom, *p* = 0.50). Histograms in upper right show the distribution of differences.

Next, we examined the model-predicted and actual fixation and perisaccadic neural responses for ensembles with RFs far from the ST. Similar to figure 4a, we transformed the axes to examine the relationship between bias-relevant STUs and the model response for ensembles with *d* ≥ 11 (Fig. 5a). Results look similar for MT and V4 neurons (Supplementary Fig. 3). Since there are two regions of bias-relevant STUs for this group of neuronal ensembles (Fig. 3a bottom), the contours in figure 5a also illustrate two temporal regions of the model-predicted response that might contribute to spatial bias. Near and far probes are defined as in figure 4b; however, for these ensembles most of the neurons have RFs on the other side of the FP from the ST (see prevalence in Fig. 5b). Since there are two regions of the model-predicted response that could potentially contribute to spatial bias, we compared the model’s response at near vs. far probes during fixation (-500:-100 ms), perisaccadic (0:40 ms), and postsaccadic (70:200 ms) windows (Fig. 5c top row). We quantified responses in a 60ms window covering the peak of each response (shown by the gray bars, fixation: 30:90 ms, perisaccadic: 40:100 ms, postsaccadic: 30:90 ms). During fixation, there was no response to either near or far probes (near = 0.99±0.00, far = 0.99±0.00, *p* = 0.86). Perisaccadically, responses were observed for both near and far probes, with a larger increase of response for near probes (near = 1.05±0.01, far = 1.01±0.01, *p* < 0.001). Postsaccadically, there was a continued increase in response at near probes, but the response at far probes decreased back to the fixation level (near = 1.06±0.01, far = 0.99±0.00, *p* < 0.001). Neuron’s responses mirror the model’s predictions in the fixation window (near = 0.98±0.01, far = 0.10±0.01), perisaccadic window (near = 1.07±0.02, far = 1.03±0.02), and postsaccadic windows (near = 1.12±0.01, far = 1.05±0.01) (Fig. 5c from top to bottom). Figure 5d demonstrates that there was no significant difference between firing rates at near vs. far probes during fixation (near = 26.27±1.01 Hz, far = 26.33±1.02 Hz, *p* = 0.31), but during the perisaccadic response window the neural firing rate for near probes was significantly higher than the firing rate for far probes (near = 28.51±1.08 Hz, far = 27.21±1.07 Hz, *p* < 0.001) and continues during the postsaccadic response window (near = 29.27±1.02 Hz, far = 27.27±1.01 Hz, *p* < 0.001). Neurons with RFs far from the ST did not respond to either near or far probes during fixation, but responded more to near probes perisaccadically and postsaccadically. Neurons responded more strongly to near-ST stimuli closer to the FP, reflecting the spatial bias opposite to the saccade direction in ensembles with *d* ≥ 11. These findings demonstrate how this systematic and unbiased search in the space of spatiotemporal sensitivity components can identify the neural basis for a biased representation of visual space during eye movements.

**Fig. 5.**
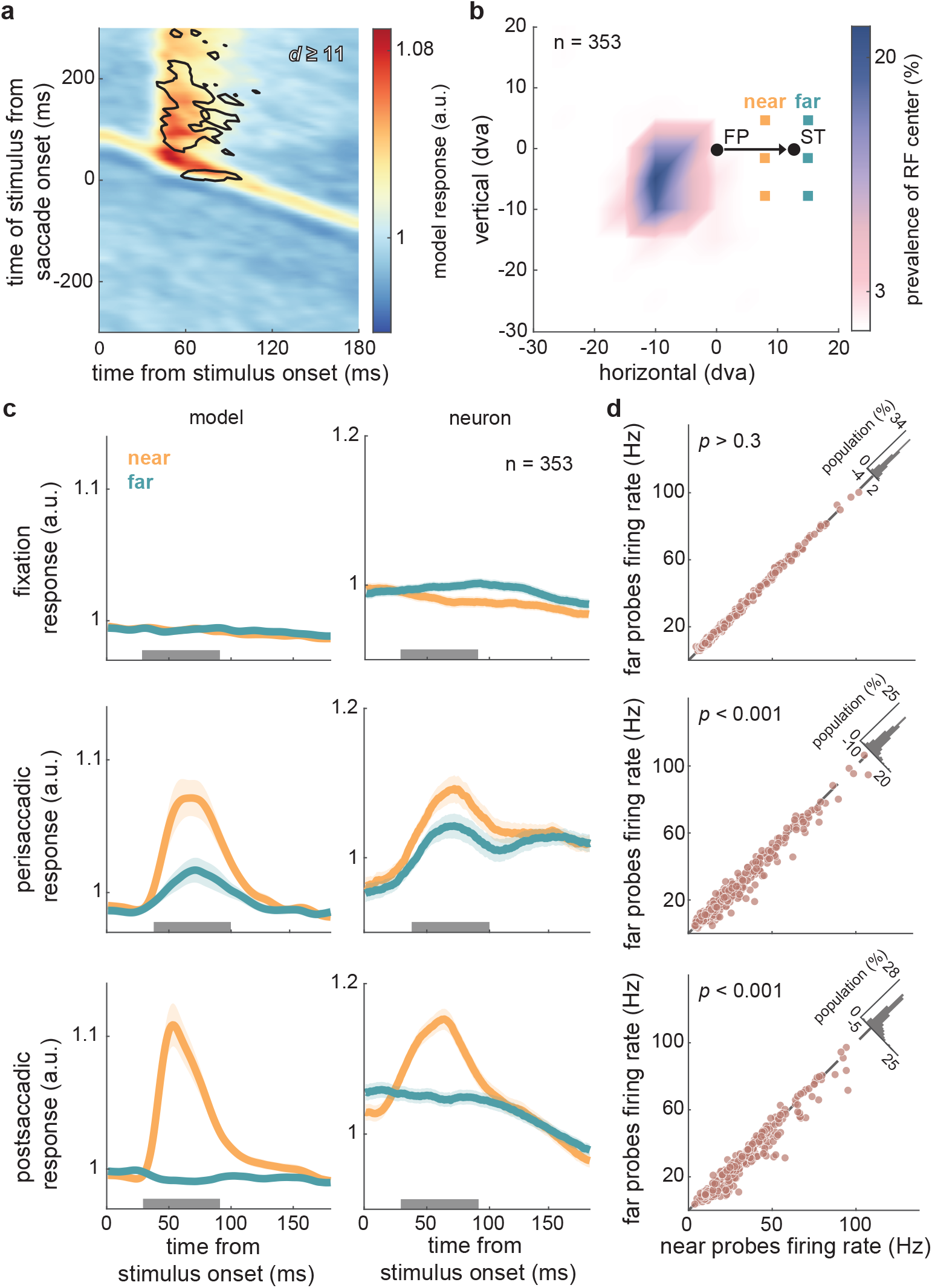
Identifying the neural correlates of spatial bias around the ST area from neurons whose RFs are located far from the ST. **a.** Bias-relevant STUs and model-predicted response, plotted as a function of time from stimulus to saccade onset (y-axis) and time of response from stimulus onset (x-axis). Shown for ensembles with *d* ≥ 11 dva (n = 11 ensembles, 353 neurons). **b.** The map of RF centers and probe locations. Similar to Fig. 4b, but for ensembles with RF centers primarily on the opposite side of the FP from the ST (defined in (a)). **c.** Mean of normalized model-predicted responses (left) and actual neural responses (right) over time from stimulus onset, for near (orange) and far (green) probes, during fixation (top; -500:-100 ms), perisaccadic (middle; -20:10 ms), and postsaccadic (60:230 ms) windows. Mean±SEM across models or neurons in the ensembles; gray bars show analysis windows used in (d). **d.** Comparison of actual neural responses to near vs. far probes (n = 94 neurons), for the fixation (top, *p* = 0.31), perisaccadic (middle, *p* = 9.44e-10), and postsaccadic (bottom, *p* = 1.23e-13) periods. Histograms in upper right show the distribution of differences.

To gain a deeper understanding of the nature of perisaccadic mislocalization, we wanted to investigate how perisaccadic neural modulations are associated with the representation around the ST and how it might be related to the observed spatial bias. To assess the change of neuronal sensitivity around the ST in the corresponding time and delay windows as the spatial bias, we defined an ST sensitivity index using kernels averaged over delays of 50:100 ms. Out of the 6 ST probes, we divided the range of kernel weights by the mean kernel weight over all times relative to saccade onset to quantify the difference in sensitivity of a neuron to various probes presented around the ST area across time from saccade onset (Fig. 6a). We excluded 95 neurons with high kernel weights during the second fixation period (240:440 ms from saccade onset) comparing to the first fixation period (-441:-241 ms) (i.e., neurons whose postsaccadic RF included the near-ST probe locations) to reduce the interference of future field activity. In the same perisaccadic time window that we observed the spatial bias (50:150 ms shown by gray bar in Fig. 6a), there was an increase in the ST sensitivity index compared with the fixation window (-300:-150 ms) (Fig. 6b; fixation = 3.27±0.07, perisaccadic = 3.67±0.09, *p* = 0.04), indicating that the modulation of neurons’ spatiotemporal sensitivity around the time of saccades enhances the representation of the ST area. To examine the relationship between the spatial bias and enhanced ST representation, we measured the ST sensitivity index again with the reduced model in which bias-relevant STUs were nullified (Fig. 3b). In the same perisaccadic window, the ST sensitivity index in the reduced model was significantly smaller than in the full model (Fig. 6c, 3.31±0.12, *p* < 0.001). Thus, the enhanced ST sensitivity index around the ST relies on the bias-relevant STUs, and a computational manipulation that removes spatial bias leads to decreased sensitivity around the ST. This reveals that the perisaccadic spatial bias could be a result of the same changes in sensitivity which enhance the ST representation around the time of saccades.

**Fig. 6.**
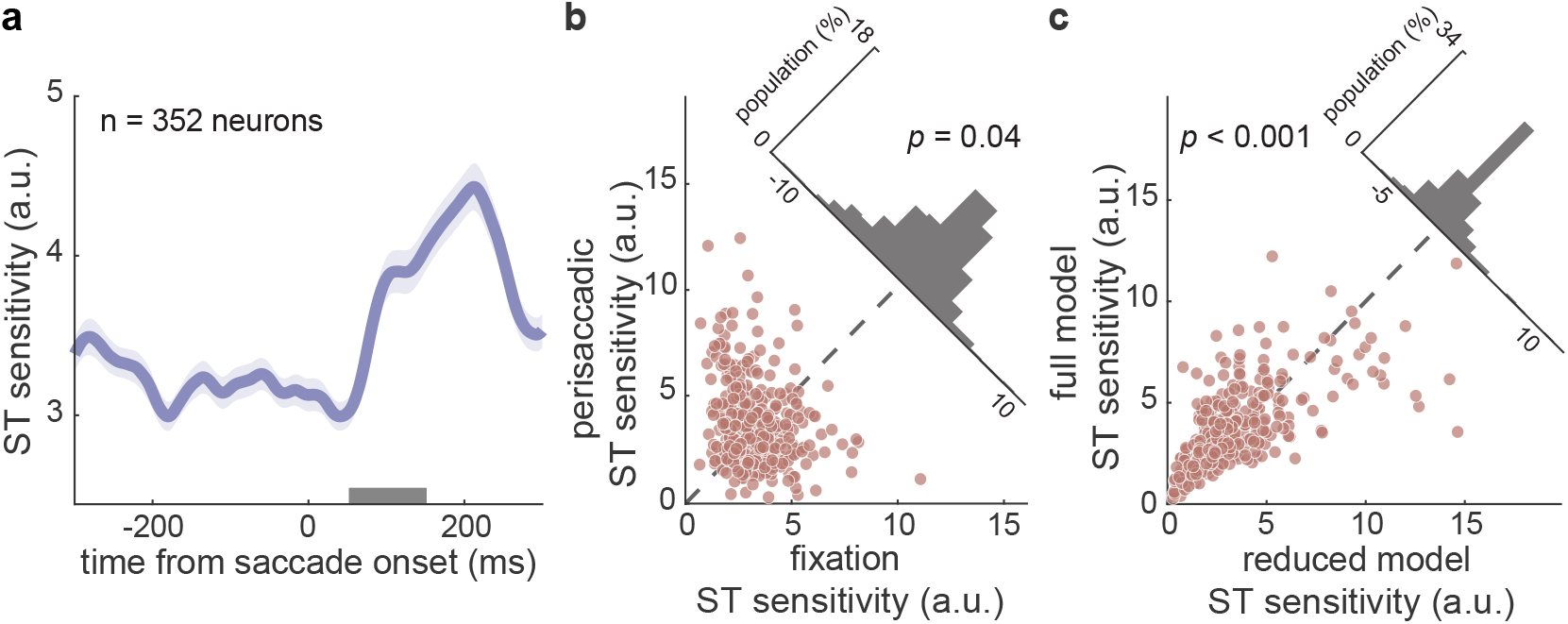
Perisaccadic enhancement of the ST representation and its relationship to the perisaccadic spatial bias. **a.** Average ST sensitivity index of 352 neurons over time from saccade onset for stimulus delay of 50:100 ms relative to each timepoint. Mean±SEM across neurons; gray bar shows the analysis window used in (b) and (c). **b.** Comparison of ST sensitivity index in the perisaccadic window (50:150 ms) and fixation window (-300:-150 ms) (*p* = 0.04). Histogram in upper right shows the distribution of differences. **c.** Comparison of ST sensitivity index in the perisaccadic window using the kernels from the full model vs. those from the reduced model (bias-relevant STUs removed) (*p* = 3.18e-09). Histogram in upper right shows the distribution of differences.

## Discussion

How the location of visual stimuli is represented in the brain is not well understood. Imaging studies have suggested that the perceived location could be encoded in extrastriate visual areas along with other visual features^54,55^. Our perception of location changes around the times of saccades^2,4^, as do extrastriate responses^27,41,56^. We used a combined experimental and computational approach to examine how changes of sensitivity in MT and V4 could explain perisaccadic mislocalization. We quantified perisaccadic spatial bias around the ST and identified the STUs relevant for the observed bias, which reveals that neurons with RFs far from the ST contribute more to the perisaccadic spatial bias. We found perisaccadic changes in extrastriate sensitivity in the identified bias-relevant time and delay windows, supporting the hypothesis that location representation occurs in extrastriate visual areas. In addition, we demonstrated that the spatial bias is accompanied by the perisaccadic enhancement of neural sensitivity around the ST, with matching time course and underlying neural response components, suggesting that the brain prioritizes saccade target representation at the expense of biases in location perception.

The existing psychophysics results have been mixed, but our neurophysiological results are consistent with many aspects of the previous literature. In total darkness, Honda reported that mislocalization in human subjects starts in the same direction as saccade and then is reversed to the opposite direction, with the greatest mislocalization occurring around 50 ms after saccade onset ^57,58^. In a double-saccade task, Jeffries et al. found that mislocalization in rhesus monkeys is in the direction opposite to the first saccade, with the maximum mislocalization around 100 ms after saccade onset^11^. Based on the model’s kernels, we observed spatial bias in the direction opposite the saccade, at a timing consistent with both the human and nonhuman primate studies (Fig. 2d); however, we cannot rule out the possibility that examining different RF or probe positions could reveal cases of spatial bias in the saccade direction. In addition to mislocalization parallel or opposite to the saccade direction, many studies have reported compression when conducting the experiments in a dimly lit room^15–17^, meaning that stimuli are perceived as closer to the saccade target (i.e., mislocalization opposite the saccade direction for stimuli past the saccade target, and in the saccade direction for others). In a computational study, Krekelberg et al. also predicted mislocalization in the direction of the saccade at a location close to the FP, and mislocalization in the opposite direction at locations near and past the ST^19^. They implemented a decoder using nonhuman primate neural data recorded from area MT, the medial superior temporal area (MST), the ventral intraparietal area (VIP), and the lateral intraparietal area (LIP) in the dark. We only found spatial bias opposite to the saccade direction for stimuli around the ST; however, we are not ruling out the possibility of a compression phenomenon, because in this study we did not measure spatial bias for stimuli at locations other than the ST (nor were our probe positions optimized to make such systematic measurements across the rest of the visual field).

Our results substantiate the association between perisaccadic mislocalization and RF remapping^19,43,44^. Like previous studies attempting to understand this connection, our approach for measuring bias assumes the same decoding algorithm is used during fixation and around the time of saccades, with altered visual responses driving the perisaccadic perceptual changes. Many studies have interpreted perisaccadic mislocalization as a flaw in the visual system while shifting the coordinate systems across saccades^13,58,59^, but it is not clear what the reason for this flaw is, or if it is the byproduct of another, beneficial, set of changes. The saccade target theory has hypothesized that the brain biases toward representing the ST in order to maintain visual stability, and the representation of non-target locations is consequently reduced^60,61^. Our results demonstrated that removal of bias-relevant neural components is correlated with a reduction of perisaccadic sensitivity around ST (Fig. 6c). Based on our results, we suggest that spatial mislocalization could be a result of allocating more neural resources toward the ST representation. The spatial bias could therefore be interpreted as a tradeoff the brain makes to amplify the ST representation perisaccadically, consistent with the saccade target theory and ST remapping. It should be noted that future field remapping could also contribute to perisaccadic spatial bias. Figure 5c shows increased perisaccadic response around the ST that might be induced by ST remapping, and the increased postsaccadic response could reflect future field remapping. This possible correlation between future field remapping and mislocalization will require further investigation. We also cannot definitively state whether these spatial biases in responses arise first in MT and V4 or are inherited from upstream areas.

Our approach in this study also reinforces the feasibility of using a GLM framework to model higher visual areas. The classical GLM has been widely used for encoding and decoding neural responses in low-level visual areas (such as the retina, LGN, and V1)^62,63^, but they fall short in capturing the time-varying characteristics of higher-level visual areas. To model responses in these areas, nonstationary model frameworks that enables a time-varying extension of a GLM have been developed, which showed success in characterizing the perisaccadic spatiotemporal changes of neural response and reading out perisaccadic stimulus information on the same timescale of saccadic eye movements^42,49,64,65^. In the present study, we took advantage of this GLM framework (SVGLM) and developed a procedure to measure spatial bias based on instantaneous neural sensitivity at various locations to identify the neural components contributing to spatial bias. Our results provide a potential explanation of the neural basis of mislocalization, which could be tested most definitively through experiments combining psychophysical measurements in macaques with causal manipulations of neural activity. These applications of the SVGLM framework demonstrate that a GLM-based approach is a viable way of studying the complex dynamics in higher-level visual areas, and could also be used to link specific aspects of neural sensitivity to different perceptual phenomena in other brain areas.

## Methods

### Behavioral paradigm and electrophysiological recording

We trained and recorded from four adult male rhesus macaques (*Macaca mulatta*). The behavioral task used in this study was a visually guided saccade task, with probe stimuli appearing at pseudorandom locations before, during, and after the saccade. To start a trial, the monkey held fixation on a central fixation point (FP). While the monkey was holding fixation, a saccade target (ST) appeared 13 dva away from the FP horizontally. In each recording session, there was only one saccade direction (leftward or rightward). After a randomized time-interval (uniform distribution between 700 and 1100 ms), the fixation point disappeared, which was the go cue for the monkey to saccade to the ST. The monkey then held fixation on the ST for 560:750 ms to receive a juice reward. Throughout the length of each trial, a complete sequence of 81 probe stimuli flashed on the screen in pseudorandom order, one at a time for 7 ms each. The probe locations were selected pseudorandomly from a 9×9 grid of possible locations. Each probe stimulus was a white square (full contrast), 0.5 by 0.5 degrees of visual angle (dva), against a black background. Each probe stimulus occurred at each time in the sequence with equal frequency across trials.

During each neurophysiological recording session, the grid of possible locations of the probes was placed and scaled to cover the estimated presaccadic and postsaccadic RF centers of the neurons recorded, the FP, and the ST. The probe grids varied in size horizontally from 24 to 48.79 (40.63 ± 5.93) dva, and vertically from 16 to 48.79 (39.78 ± 7.81) dva. The distance between two adjacent probe locations varied horizontally from 3 to 6.1 (5.07 ± 0.74) dva, and vertically from 2 to 6.1 (4.97 ± 0.97) dva.

While the monkey was performing the task, we monitored their eye movements with an infrared optical eye-tracking system (EyeLink 1000 Plus Eye Tracker, SR Research Ltd., Ottawa, CA) with a resolution of <0.01 dva (based on the manufacturer’s technical specifications), and a sampling frequency of 2 kHz. Presentation of the visual stimuli on the screen was controlled using the MonkeyLogic toolbox. In total, 332 neurons in the middle temporal (MT) cortex and 291 neurons in area V4 were recorded in 108 sessions, but only 300 MT and 147 V4 neurons were used in order to make ensembles of neurons with at least 10 neurons with a similar RF, ST and grid position during recording. We recorded both spiking activity and the local field potential (LFP) from either MT or V4 using 16-channel linear array electrodes (V-probe, Plexon Inc., Dallas, TX; Central software v7.0.6 in Blackrock acquisition system and Cheetah v5.7.4 in Neuralynx acquisition systems) at a sampling rate of 32 KHz, and sorted neural waveforms offline using the Plexon offline spike sorter and Blackrock Offline Spike Sorter (BOSS) software.

### RF center estimation

The centers of RFs were assigned based on responses to the probes that generated the maximum firing rate during the fixation period before the saccade. For each probe location, the probe-aligned responses are calculated by averaging the spike trains over repetitions of the probe before or after the saccade (greater than 100 ms before or after the saccade onset), from 0:200 ms following probe presentation, across all trials. The response is then smoothed using a Gaussian window of 5 ms full width at half maximum.

### Encoding model framework

The Sparse Variable Generalized Linear Model (SVGLM) used in this study was previously developed by Niknam et al.^53^, see this paper for more details of the model fitting. The SVGLM is a variant of the widely used GLM framework^62,63,66^ that tracks the fast dynamics of sparse spiking activity with high temporal precision and accuracy. The SVGLM is a model that captures the neurons’ sensitivity varying over space and time with high temporal resolution by using a reduced number of STUs selected through a dimensionality reduction process (see Supplementary Information). The fitted model also captures how much these STUs contribute quantitatively to generating spikes on a precise millisecond timescale during a saccade. The weighted combination of these STUs constitutes the spatiotemporal stimulus kernels. The SVGLM defines a conditional intensity function according to the equation,

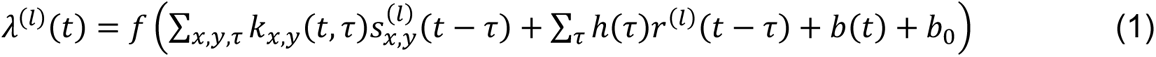

where 𝜆 is the instantaneous firing rate of the neuron at time 𝑡 in trial 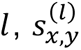 is either 0 or 1 representing respectively the off or on condition in a sequence of probe stimuli presented on the screen at probe location (𝑥, 𝑦) in trial 𝑙. 𝑟(𝑙)(𝑡) denotes the spiking response of the neuron for trial 𝑙 and time 𝑡, 𝑘_𝑥,𝑦_(𝑡, 𝜏) represent the stimulus kernels, ℎ(𝜏) indicates the post-spike kernel applied to the spike history which captures the refractory effect, 𝑏(𝑡) is the offset kernel that reflects the change of baseline activity induced by saccades, the constant 𝑏_0_ = 𝑓^−1^(𝑟_0_) with 𝑟_0_ as the measured mean firing rate (Hz) across all trials in the experimental session, 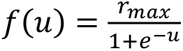 is a static sigmoidal function that describes the nonlinear properties of spike generation with 𝑟*_max_* indicating the maximum firing rate of the neuron obtained empirically from the experimental data. The model was fitted using an optimization procedure in the point process maximum likelihood estimation framework^67^ at the level of single trials. The evaluation for model performance is described in supplementary information (Supplementary Fig. 1 b-d).

### Measuring spatial bias

Neurons recorded with the same ST position and probe arrangements (grid positioning and spacing), and with similar RF locations were grouped as an ensemble. 15 ensembles were formed, each with a minimum of 10 neurons. Before any analysis, kernels of all neurons were smoothed by moving average with time windows of 50 ms for time 𝑡 and 20 ms for delay 𝜏 to reduce noise. Figure 1c shows 9 kernels for a sample probe at the center of a neuron’s RF and its 8 neighboring probes, stacked over neurons in an example ensemble. For each particular time and delay, we constructed two population kernel vectors consisting of the kernel values of center probe and a neighbor probe at that time and delay with all neurons in an ensemble. To measure the similarity between kernels at neighboring probe locations, we computed the correlation between the kernels of center probe and a neighbor probe with all neurons in an ensemble, and subtracted baseline correlation values during fixation (-441:-141 ms from saccade onset). The correlation was measured for each of the 8 neighboring probes, and repeated for 7×7 probe locations (after excluding probes on the edges). Using correlation values as magnitudes, and the probe position relative to the center probe as directions, we formed 8 vectors at each probe location across time and delay, and took the average of these 8 vectors at each of the 7×7 probe locations. The polar plot in Fig. 1e shows these vectors between a sample probe around ST and its 8 neighboring probes. Spatial bias at each location was defined as the horizontal projection of the average vector at that location, and it was computed for all 15 ensembles. These spatial bias values were used to construct spatial bias maps across time and delay for each of 7×7 probe locations. Figure 2a-b shows two cross-sections of an example bias map, associated with a sample probe location around ST, over particular time and delay windows. For each ensemble, we averaged the bias maps at the 6 probe locations closest to the ST, excluding probes that were within 2 dva from either the presaccadic or postsaccadic RF (Fig. 2c). Before averaging the bias maps of all 15 ensembles, the spatial bias of each ensemble was normalized to range from -1 to 1. We used one-sided Wilcoxon signed-rank test to report *p*-values for all our statistical comparison analysis, if not mentioned specifically.

### Identifying modulated STUs

To identify which components of the neurons’ spatiotemporal sensitivity drive the neuron’s response changes around the time of saccades, we first quantify the contribution of each STU. It is expected that out of all STUs, only some of them at specific times and delays contribute to the generation of the neural response (referred to as ‘contributing STUs’). These contributing STUs are identified during the dimensionality reduction process during the model fitting, based on a statistically significant contribution to the stimulus-response relationship (see Niknam et al.^53^ for details).

We then define the modulated STUs as those for which the fraction of contributing STUs in a 3×3 window around that STU’s time and delay is significantly different during perisaccadic period compared to fixation period. Mathematically speaking, the fraction of contributing STUs needs to fulfill the following condition:

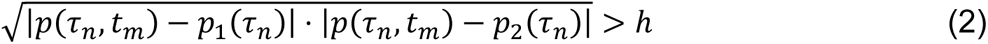

with 𝑝(𝜏_𝑛_, 𝑡_𝑚_) as the fraction of contributing STUs in a 3×3 window around the 𝑛 ^th^ bin of delay and 𝑚 ^th^ bin in time 1 < 𝑛 < 30,1 < 𝑚 < 156. 𝑝_1_(𝜏_𝑛_) is the fraction of contributing STUs during the first fixation period 540 to 120 ms before saccade in time bin 1 to 60 at 𝑛 ^th^ bin of delay (1 < 𝑛 < 30), and 𝑝_2_(𝜏_𝑛_) is the fraction of contributing STUs during the second fixation period 280 to 540 ms after saccade in time bin 120 to 156 at 𝑛 ^th^ bin of delay (1 < 𝑛 < 30). ℎ is a significance threshold between 0 and 1, and was set to 0.3 for the analysis.

### Identifying bias-relevant STUs

From the list of modulated STUs, we identified the ones that contribute to spatial bias specifically (termed bias-relevant STUs). The contribution of each modulated STU to the spatial bias was quantified by evaluating its impact on the difference between kernels at neighbor probes across a saccade, by removing each modulated STU one at a time and testing if the change in kernels difference is significant based on a bias index. Because spatial bias was measured from the correlations between kernels at neighbor probes for an ensemble of neurons, the difference between the stimulus kernels of two neighboring probes for individual neurons, may impact the resulting spatial bias read out from that ensemble. The absolute difference between each pair of stimulus kernels of the fitted models at two neighboring probe locations (𝑥_0_, 𝑦_0_) and (𝑥_𝑖_, 𝑦_𝑖_) and at each delay (𝜏) across different times to the saccade (𝑡), was quantified as

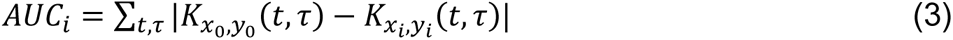

where 𝐴*UC*_𝑖_, represents the area under curve of the difference of kernels 𝐾_𝑥0,𝑦0_ (𝑡, 𝜏) and 𝐾_𝑥𝑖,𝑦𝑖_ (𝑡, 𝜏) over time and delay, between each center probe at (𝑥_0_, 𝑦_0_) and each of the eight neighbor probes (𝑖 ∈ {1, …, 8}) at (𝑥_𝑖_, 𝑦_𝑖_) (Supplementary Fig. 2 A). Since the spatial bias was measured for 6 probe locations around ST, we also measured the difference AUCs at those same 6 center probes for the neurons in each ensemble. The average of difference AUCs over 8 center-surround probe pairs was used to compute the bias index associated with individual center probe in the following. For each neuron in each ensemble, we first measured the difference AUCs with the full model (no perturbation in the model estimated STUs). Next, we remove each of the modulated STUs one at a time from the full model by replacing that STU with zero and repeat the above steps so that we have a list of difference AUCs measured without each of the modulated STUs. We then define the bias index for each modulated STU as the absolute difference between the AUC for full model and the AUC corresponding to removing each of the modulated STU from the model (Supplementary Fig. 2b). To identify bias-relevant STUs, we define a threshold for this bias index as the 90^th^ percentile of the cumulative distribution function of all the nonzero bias indices (bias index of 2.56) as the threshold (Supplementary Fig. 2c). A larger bias index means that nulling the weight of a particular STU results in a stronger change in kernel differences for the probes around the ST, so the STUs with a bias index above the threshold were classified as bias-relevant. The bias indices were specific to each of the 6 ST probes and for each neuron. Figure 3a shows the mean bias index across probes and neurons which is used to generate the map of bias-relevant STUs. To validate if the identified bias-relevant STUs using this procedure would actually contribute to the readout spatial bias from each ensemble, we removed the identified bias-relevant STUs from the model for each neuron, and recomputed the spatial bias (Fig. 3b).

## Data and code availability

The datasets generated and/or analyzed for this study will be available on a public repository upon the acceptance of the manuscript.

## Acknowledgements

The authors would like to thank the animal care personnel at the University of Utah. We specifically thank Rochelle D. Moore and Dr. Tyler Davis for their assistance with the NHP experiments, and Dr. Kelsey Clark for her comments on the manuscript. This work was supported by NIH EY026924 & NIH NS113073 to B.N.; NIH EY031477 to N.N.; NIH EY014800 and an Unrestricted Grant from Research to Prevent Blindness, New York, NY, to the Department of Ophthalmology & Visual Sciences, University of Utah.

## Author contributions

N.N. and B.N. conceived the study. B.N. performed the surgical procedures. B.N., N.N., and A.A. designed the experiment. B.N., N.N., K.C., and G.W. designed the analysis. B.N. and A.A. performed the physiology experiments and acquired data. A.A. performed the modeling. G.W. performed the data analysis. G.W., K.C., N.N., and B.N. wrote the manuscript.

## Competing interests

The authors declare no competing interests.

**Supplementary Information** is available for this paper.

## Notes

### Competing Interest Statement

The authors have declared no competing interest.

