## Supplementary Information for "Neural correlates of perisaccadic visual mislocalization in extrastriate cortex"

Behrad Noudoost

Neda Nategh

**Word counts**

1030

**Number of figures**

3

**Dimensionality reduction**

The fast dynamics of the neurons' spatiotemporal sensitivity around saccades requires a high-dimensional representation of STUs. We designed the behavioral paradigm with probe stimuli presenting every 7 ms to establish a set of temporal basis functions $\mathcal{B}_{i,j}(t,\tau)$ that down-sample the time $t$ and delay $\tau$ into 7-ms bins using second order B-spline functions $\mathcal{U}_i(\tau)$ and $\mathcal{V}_j(t)$:

$$\mathcal{B}_{i,j}(t,\tau) = \mathcal{U}_i(\tau)\mathcal{V}_j(t) \quad (1)$$

$\{\mathcal{U}_i(\tau)\}$ spans across delay $\tau$ and represents a kernel that lasts for 200 ms, constructed by a set of 33 evenly spaced knots at {-13, 6, …, 204, 211} ms, which correspond to a total of 30 basis functions. Similarly, $\{\mathcal{V}_j(t)\}$ spans across time $t$ and represents a saccade-aligned kernel that lasts for 1081 ms, constructed by a set of 159 evenly spaced knots at {-554, -547, …, 545, 552} milliseconds, which correspond to a total of 156 basis functions.

Binning the time and delay dimensions reduces the dimensionality by about two orders of magnitude, which is still not feasible for a computationally robust estimation. To limit the amount of STUs in the estimation, we used a statistical method to identify those STUs that have a significant impact on a neuron's response at a specific time[1]. We compared the weight distribution of the STU by fitting a generalized linear model (GLM) on 100 subsets of randomly selected spike trains (35% of all trials) to a control distribution obtained from 100 subsets of shuffled trials where the relationship between stimulus and response was altered. The conditional intensity function (CIF) of this GLM is defined as:

$$\lambda_t = f\left(\sum_{\tau=1}^{T} S_{x,y}(t-\tau)\cdot\kappa\cdot\mathcal{B}_{i,j}(t,\tau)\right) \quad (2)$$

with $\lambda$ as the instantaneous firing rate of the neuron, $S$ is the stimulus history of length $T$ at location $(x,y)$, $\kappa$ is the weight of a single STU, represented by basis function $\mathcal{B}_{i,j}(t,\tau)$, whose significance of contribution is evaluated by satisfying the following condition:

$$|\mu - \tilde{\mu}| \geq 1.5\tilde{\sigma} \quad (3)$$

which denotes that the absolute difference between the mean of the original weight distribution $\mu$ and the mean of the control weights distribution $\tilde{\mu}$ should be above or equal to 1.5 times the standard deviation of the control weights distribution $\tilde{\sigma}$. The threshold of 1.5 was chosen heuristically to reduce the dimensionality of STU space to ~$10^4$, making the model fitting process practical without overfitting. Next, we use this subset of STUs to parameterize the linear filtering stage of an encoding model in a less complex space, with the aim of determining how these STUs build up the neuron's spatiotemporal sensitivity map. The STUs' weighted combination over time $t$, delay $\tau$, and probe $(x, y)$ describes the neuron's sensitivity kernels $k_{x,y}$ at each time point relative to saccade onset as below:

$$k_{x,y}(t,\tau) = \sum_{i,j} \kappa_{x,y,i,j} \cdot \mathcal{B}_{i,j}(t,\tau) \quad (4)$$

where $\{\kappa\}$ are the weights of the STUs obtained from the encoding model (defined in Eq. (1) in Methods). Note that the sum is limited to the subset of $\mathcal{B}$ whose corresponding STU was considered significant based on Eq. (3), and the weights for the remaining STUs were assigned a value of zero.

**Model performance**

The data were randomly split into a training set (35%), a validation set (30%), and a testing set (35%), and we used the testing set to evaluate the model performance. Supplementary figure 1b shows that the model predicts the neural response at the single trial level. The “good” trial was defined as the trial with the largest normalized log-likelihood difference (ΔLL), and the “average” trial has the median normalized ΔLL. The normalized ΔLL was calculated as the log-likelihood (LL) of the spike trains using the model-predicted firing rate minus that under a null model and normalized by spike counts. The null model is a model where the instantaneous firing rate of the neuron is set to its average firing rate. Supplementary figure 1c compares the normalized ΔLL for fixation (-300:-150 ms) vs. perisaccadic (0:150 ms) time windows and shows that the performance of model-predictions in the perisaccadic period is slightly better than in fixation period (fixation = 0.15±0.00 bits/spike, perisaccadic = 0.16±0.00 bits/spike, $p$ = 0.00), indicating that the model is successfully capturing changes in neural sensitivity around the time of saccades. Supplementary figure 1d shows the correlation coefficient (CC) between the model-predicted firing rate and the empirical firing rate in response to the repeated presentation of a sequence of probe stimuli falls within the level of the inherent trial-by-trial variability. The data-data CC was measured between binned firing rates in response to the same 300 ms stimulus sequence; data were randomly split (60%-40%) 15 times and the mean is reported. The average firing rate was computed by binning the probe-aligned spikes using non-overlapping windows of 30 ms and smoothing the binned response with a Gaussian window of 5 ms (full width half max) and normalizing to have a mean of zero and unit standard deviation. The data-data CC is significantly higher than the model-data CC (data-data = 0.44±0.01, model-data = 0.30±0.01, $p$ = 5.96e-75).

*Comparing responses and bias-relevant STP maps between MT vs. V4 neurons*

Supplementary figure 3 shows the model response for MT vs. V4 neurons for neurons with $d$ < 11 and $d \geq 11$. For MT neurons with $d$ < 11, we outlined the regions corresponding to bias-relevant STUs with 60% contour, which is around time from stimulus onset 70:110 ms and time of stimulus from saccade onset -20:10 ms (top left). V4 neurons with $d$ < 11 was outlined with 50% contour and show bias-relevant response at a slightly earlier around time from stimulus onset 60:100 ms and time of stimulus from saccade onset -20:10 ms (top right). MT neurons with $d \geq 11$ have two regions of bias-relevant response at 52% contour (bottom left). The first region is around time from stimulus onset 60:100 ms and time of stimulus from saccade onset 0:20 ms, and the second region is around time from stimulus onset 40:100 ms and time of stimulus from saccade onset 40:230 ms. Similarly, V4 neurons with $d \geq 11$ was outlined with 56% contour, and the first region of bias-relevant response is around time from stimulus onset 50:100 ms and time of stimulus from saccade onset 0:20 ms, and the second region is around time from stimulus onset 40:100 ms and time of stimulus from saccade onset 70:240 ms (bottom right).

### Supplementary figures and figure legends

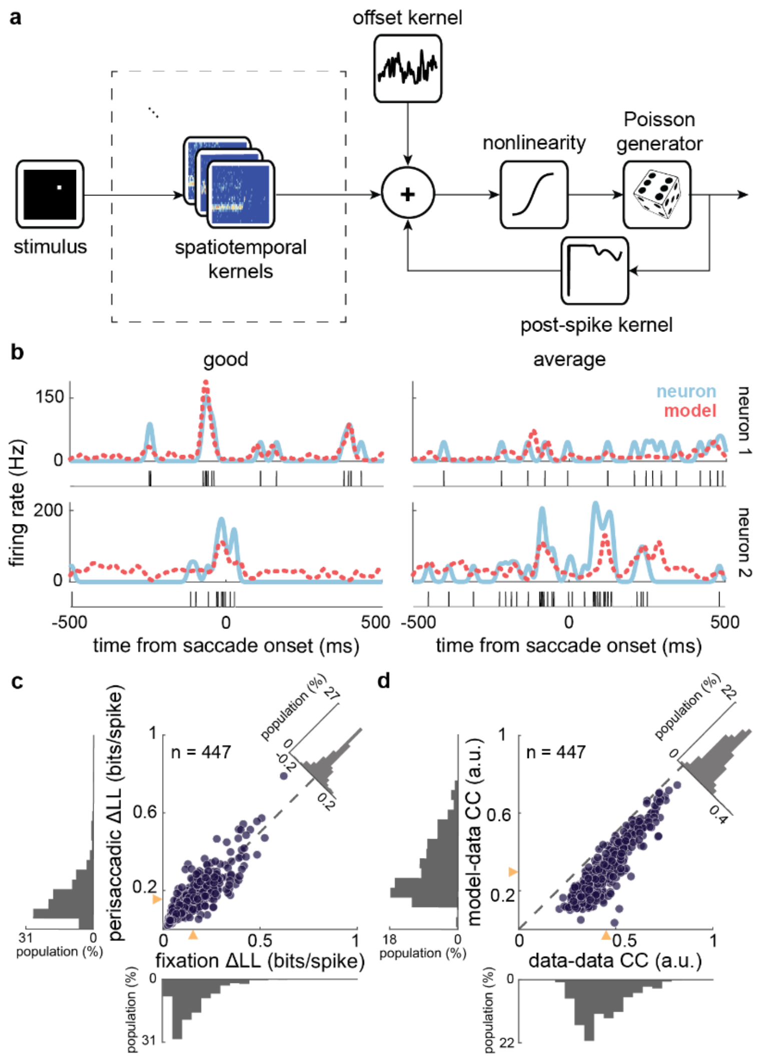


**Supplementary Fig. 1. Structure and performance of the SVGLM. a.** Schematic of the SVGLM. The stimulus is convolved with 4-dimensional kernels (consisting of STUs) representing the time-varying spatiotemporal sensitivity of individual neurons. The filter stimulus is then added to the output of an offset kernel and the signal generated by a post-spike kernel. The sum passes through a nonlinearity to estimate the neuron’s spike rate which is used as the underlying rate by a Poisson spike generator to predict the neuron’s spiking activity. **b.** The recorded neural response vs. model-predicted response for a “good” trial (best ΔLL) and an “average” trial (median ΔLL) of two example neurons. Spikes in each trial are shown below the smoothed traces. **c.**

Comparison of the normalized ΔLL of the recorded spikes under the model-predicted response in fixation (-300:-150 ms) vs. perisaccadic (0:150 ms) windows. Yellow triangles illustrate the medians (fixation = 0.15, perisaccadic = 0.16); histograms show the marginal distributions (left, bottom) and the difference distribution (upper right). **d.** Comparison of the normalized CC between the data-data correlation vs. model-data correlation, evaluating variability in responses to the same stimulus. Yellow triangles illustrate the medians (data-data = 0.45, model-data = 0.30); histograms show the marginal distributions (left, bottom) and the difference distribution (upper right).

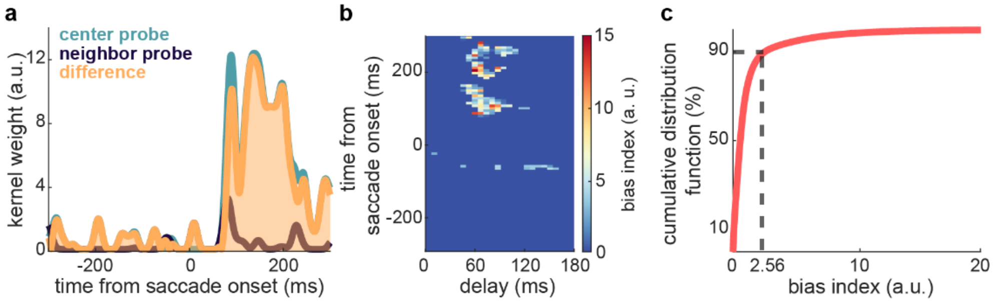


**Supplementary Fig. 2. Identifying bias-relevant STUs. a.** Pictured are example kernels for two example probe locations over 1:200 ms delay, and the difference between them, over time. The AUC represents the dissimilarity between kernels at two neighboring probes over time and delay. The process is repeated for STUs at 6 probe locations around the ST for all time and delay. **b.** Shows the map of bias index over STUs measured using difference between the full model AUC and the AUC with that STU removed. **c.** The cumulative distribution function of all the non-zero bias indices. Using the $90^{th}$ percentile as a threshold, the STUs with an absolute bias index difference above 2.56 are defined as bias-relevant.

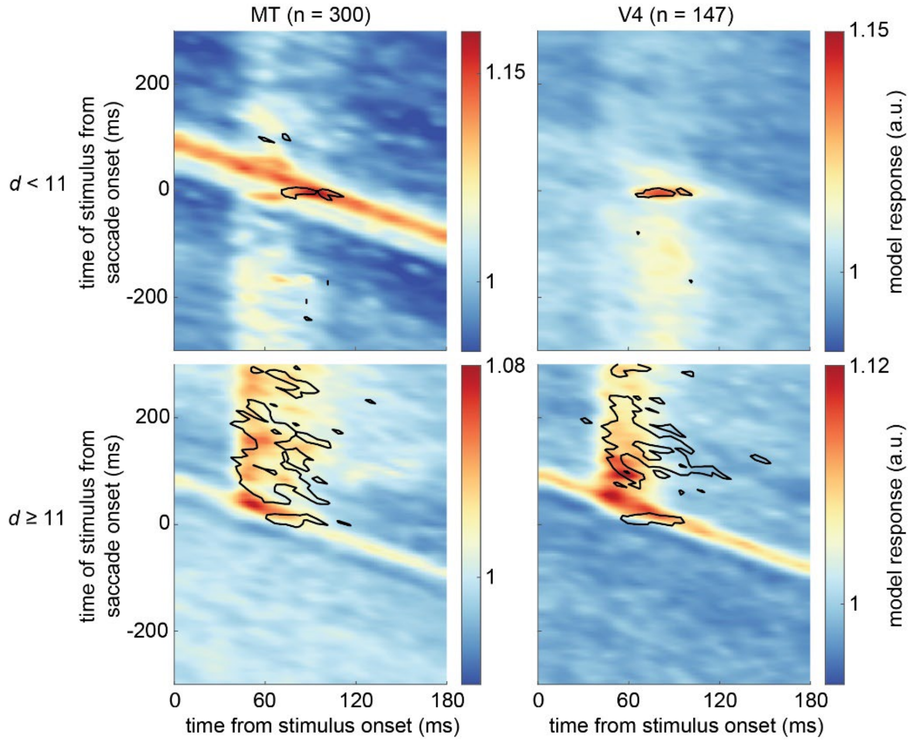


**Supplementary Fig. 3. Model-predicted responses and bias-relevant STUs for MT and V4 neurons with RFs near or far from the ST.** Color indicates model-predicted response, and black contours outline bias-relevant STUs, over time between stimulus presentation and saccade onset (y-axis) and time of response from stimulus onset (x-axis), for models of neurons recorded from MT (left) and V4 (right), for neurons with RFs near the ST (top), or far from the ST (bottom); *d* indicates the distance between the neurons' RF center and ST in dva.
